# The role of relatedness within a neighborhood in plant-to-plant interaction: effect on height growth and rust damage in *Populus nigra*

**DOI:** 10.1101/2023.06.21.545987

**Authors:** Mathieu Tiret, Leopoldo Sánchez, Martin Lascoux

## Abstract

The competitive ability of domesticated plants, which may have conferred a fitness advantage in the wild, may result in a reduction of yield in agricultural and forestry contexts, as what matters is the group rather than the individual performance. Traits related to competitive ability can be affected by the presence or absence of related individuals in their neighborhood. Consequently, local relatedness might reveal plant-to-plant interaction that can enhance the predictive abilities of genomic models when accounted for, though it remains difficult to measure. To overcome this difficulty, we analyzed data from the French breeding program of *Populus nigra* L., where 1,452 genotypes were replicated six to eight times, each time encountering a different neighborhood. We assessed local relatedness and investigated genomic estimated breeding values on tree height and vulnerability to rust with a single-step GBLUP incorporating local relatedness as a covariate. The results indicate that incorporating local relatedness as an additional factor in GBLUP models has a significantly greater influence on resistance to rust than on tree height, though its overall effect on genomic predictions themselves was limited. The influence of local relatedness is small but likely trait-specific, and the genetic architecture of the trait under selection could attenuate or improve the efficacy of breeding for group performance.

## Introduction

Breeding generally consists in redirecting evolutionary trade-offs towards new equilibria that often decrease fitness in favor of other economically interesting traits, or towards paths that were insufficiently explored by evolution in natural populations, such as group selection (Weiner, 2019). The importance of group selection, defined as the cooperation within groups or populations that would confer a selective advantage (Leigh, 2010), is controversial in nature (Maynard Smith, 1964; Lewontin, 1970; Wilson & Wilson, 2007; Okasha, 2020). However, in a breeding program, selecting for individuals that improve the overall performance (e.g., robust productivity, tolerance to biotic or abiotic stress, or average yield), by limiting unfavourable plant-to-plant interactions can be interesting (also known as ‘communal ideotype’; Biernaskie, 2022).

Separating an individual’s performance into what is due to its own genetics and to its neighbors’ has a long history of theoretical development, especially in quantitative genetics (Griffing, 1967; Bijma & Wade, 2008; Ellen et al., 2016; Bailey et al., 2018). The performance of an individual is decomposed into direct and indirect breeding values (also called direct and indirect genetic effects or DGE/IGE), where the former is the intrinsic value of an individual without any competitors, and the latter the ability of the individual to increase or decrease performance due to competition. The estimation of the direct and indirect values is based on the variation of the neighborhood of a focal individual, at the cost of a quadratic increase in data requirements. Recent theoretical developments showed that accounting for IGE can limit the detrimental competitiveness of individuals (Montazeaud et al., 2020; Bourke et al., 2021; Lemoine et al., 2023).

Breeding can especially benefit from the IGE framework when intraspecific competition is strong among elite candidates (Weiner et al., 2017). Indeed, selfish strategies in the sense of selfish genes, promoting individual fitness at the expense of the population’s collective interest (Dawkins, 1976), are often costly for the populations in breeding schemes - and breeding circumventing such strategies ended up with very successful results (e.g., Donald, 1981; Donald and Hamblin, 1983; Weiner et al., 2010). Competitive behavior can, therefore, be a burden that breeders might want to select against. Nonetheless, despite the early success of some ideotypes optimizing group performance, breeding programs do not necessarily focus on integrating this dimension (e.g., Denison et al., 2003; Murphy et al., 2017; Montazeaud et al., 2020). Consequently, the pool of elite genotypes used nowadays might be mostly composed of selfish genotypes, since early mass selection, bypassing group dynamics, favored selfish genotypes for their vigorous phenotypes (Murphy et al., 2017). It is therefore increasingly important in order to reach sustained genetic gain in breeding programs to counteract the effect of natural selection acting at the deployment phase, and to correct past artificial selection trajectories focussed solely on competitive (i.e., selfish) individuals.

In the specific case of plant breeding, resource competition between candidates occurs with the nearest plants (Casper et al., 1997; Milbau et al., 2007; File et al., 2012). The intensity of competition among neighbors may depend on their relatedness, or local relatedness (average relatedness in the vicinity of a given individual), either because of kin selection (explaining cooperation within a group because of relatedness; Hamilton, 1964; File et al., 2012; Ehlers & Bilde, 2019), or niche partitioning (explaining, on the contrary, that similar individuals may compete for similar resources; Silvertown, 2004; Guisan et al., 2005; Bengtsson et al., 2019). Local relatedness, which usually varies across trials due to varying neighborhood, can create spatial heterogeneity that, when not accounted for, biases genetic estimates. Therefore, plant-to-plant interactions can partly be due to local relatedness in one way or the other (Cahill et al., 2011), so that BLUP and derived models can be improved by incorporating the information of neighboring genotypes. Although kin recognition was reported to be not particularly relevant in crop species because of their already high relatedness (e.g., Murphy et al., 2017), we might be able to detect it in breeding schemes that have been less intensive, such as forest tree breeding with shorter linkage disequilibrium and starting with a much wider genetic base (Grattapaglia & Resende, 2011). Group selection is in fact expected to give better genetic progress for perennial species, since (i) inbreeding coefficient is much lower than in crop species (even though inbreeding depression can be higher; Lesaffre & Billiard, 2021), and (ii) positive or negative interactions between neighbors last a lifetime.

Black poplar (*Populus nigra* L.) is a Eurasian riparian forest tree that contributes as a parent, along with *Populus deltoides* Bartr. ex Marsh., to one of the most widely used hybrid tree in forest breeding (*Populus x canadensis*), and is widely deployed as clones, constituting a powerful case study of kin recognition. The dataset, stemming from the French breeding program with 1,452 genotypes replicated six to eight times, created different neighborhoods for each replicated genotype (Pégard et al., 2020), all resulting from controlled crosses of 34 parents from natural populations, some of which were already used in the breeding programme for their performance. We focused on two quantitative traits relevant for breeding, tree height and vulnerability to rust. Growth of Black poplar is particularly susceptible to foliar rust, caused by the fungus *Melampsora larici-populina* Kleb.. In order to assess the contribution of local relatedness in the context of genomic selection (Meuwissen et al., 2001), we performed genomic evaluations with a multitrait single-step Genomic Best Linear Unbiased Predictor (ssGBLUP; Legarra et al., 2009; Christensen et al., 2010). Estimating breeding values with ssGBLUP is a solid baseline in forest trees as recently shown (e.g., Cappa et al., 2019; Ratcliffe et al., 2017), and has successfully predicted growth and wood quality for many species (Lenz et al., 2020), such as Eucalyptus (e.g. Resende et al., 2012), pines (e.g., Isik et al., 2016), or spruces (e.g., Beaulieu et al., 2014).

We propose to assess the indirect genetic effect with local relatedness. More specifically, the objective of the study was to measure the extent to which local relatedness impacts the focal individuals’ phenotypes, and to quantify the benefit of including local relatedness in genomic prediction models. We also investigated the level of association between local relatedness and the traits: rust development is influenced by susceptibility of nearby trees (Pinon & Frey, 1997; Burdon & Thrall, 1999), hence is expected to be associated to local relatedness, whereas one-year-old tree height is expected to be a poor indicator of competition (as seedlings are too small to overshadow their neighbors), and is most likely related to the environmental quality of the trial. We show that (i) as expected, incorporating local relatedness as an additional factor in GBLUP models has a significantly greater influence on estimates of resistance to rust than on tree height, but (ii) its overall effect on genomic predictions is limited. Differences in traits genetic architecture and its role in plant-to-plant interactions is discussed.

## Material and Methods

### Plant material

The study is based on previously published data (Pégard et al., 2019, 2020). Seventeen male and 17 female *Populus nigra* were sampled in France, from 21 natural populations with no a priori selection and 13 from the French poplar breeding program for their performance on a range of traits, including growth and resistance to rust. These individuals were used as parents in a factorial crossing plan and double-pair mating scheme. Clones of both parents and progeny, and a hundred additional individuals from breeding programs, were grown in four sequential experimental trials (parallel strips of two lines of trees, with an intra-spacing of 1m and an inter-spacing of 2 metres) at the same location (Guéméné-Penfao, France, 47°37’59”N, 1°49’59”W), each with a randomized incomplete block design of single tree-plots (1.0 m x 2.0 m) with six blocks each, except for one trial with eight blocks. In total, the experiment comprised 10,301 trees (including 7,169 trees with genotype information), 42 full-sib families, 5 half-sib families, 1,452 genotypes (each replicated on average 7.09 times), and an Unknown Parent Group (UPG) of 105 genotypes (Table S1). Family size ranged from 1 to 119, with an average of 30.2 genotypes per family. Twenty genotypes and 17 families were shared across the four trials.

### Phenotype measurements

We focused on two phenotypic traits on one-year-old trees: height and vulnerability to rust (both measured on field). Height was assessed with a graduation rod (in cm), and vulnerability to rust (natural infection) was assessed on a scale of 1 to 9 : 1 when no rust was observed on the tree and 9 when more than 75% of lamina was covered by rust on more than 25% of the leaves (Legionnet et al., 1999). The homogeneity of rust pressure across the trial was assured by the level of infection measured on control individuals, being equally infected across blocks (the control cv. BDG that is infected with an average score of 4.37 ± 0.65 across trials, with scores ranging from 3 to 5). Some missing values were present across the trials (dead trees or unexploitable data), but at least 95.29%, 96.05%, and 95.13% of the trees were phenotyped for height, vulnerability to rust, or both traits, respectively. Given the low missing rate, we assumed that missing values only marginally unbalanced the block design, and so we used default parameters of the software packages for handling missing values when analyzing the data.

In each trial, every phenotype was first corrected for micro-environmental heterogeneity with a multitrait mixed model as follows:

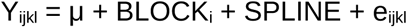

where ‘Y_ijkl_’ is the l-th phenotype of the k-th replicate of the j-th genotype in the i-th trial, μ is the grand mean, ‘BLOCK_i_’ the random effect of the i-th block, ‘SPLINE’ B-spline spatial autocorrelation correction over the surface of rows and columns (Cappa & Cantet, 2007), and ‘e_ijkl_’ the residuals. Both the block effect and the residuals follow a Gaussian distribution with diagonal covariance matrices, and the model was fitted separately by trial. We used the function *remlf90* (R package *breedR* v0.12; Munoz and Sanchez, 2014) with the parameters ‘model’ set to ‘splines’ and ‘method’ set to ‘em’. All further analyses were performed on the residuals of this model, i.e., on spatially adjusted phenotypes.

### Genomic data

Out of the 1,452 genotypes, 1,034 were genotyped (25 parents and 1,009 offspring), using a *Populus nigra* 12K custom Infinium Bead-Chip (Illumina, San Diego, CA) (Faivre-Rampant et al., 2016). Details of the DNA extraction protocol, bioinformatic pipelines, and SNP mapping on the 19 chromosomes are given in Faivre-Rampant et al. (2016) and Pégard et al. (2019, 2020). It is worth noting that the SNP chip array included SNP in QTLs and expression candidate genes associated with, among other traits, resistance to rust, but not with vertical growth. We retained SNPs with a Minor Allele Frequency (MAF) > 5%; in total, 7,129 SNPs were retained for further analyses (out of 7,513 initial SNPs).

### Genomic relationship matrix

In order to build the relationship matrix necessary for inferring breeding values, pedigree and genomic information were used as specified in the framework of the single steps estimation method (Legarra et al., 2009; Christensen et al., 2010). The pedigree relationship matrix A^Γ^ (following the notation of Legarra et al., 2009) was built using the tabular rule method (Emik and Terrill, 1949), modified to account for a single metafounder (assuming a single base population for the Unkown Parent Group or UPG; Legarra et al., 2015), with a self-relationship of γ = 8σ_p_ ^2^, with σ_p_ ^2^ the variance of allele frequencies across markers (Garcia-Baccino et al., 2017). The combined relationship matrix H was summarized in Aguilar et al. (2019), and is:

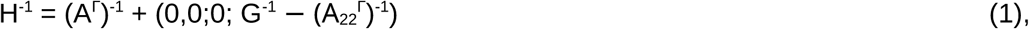

where A ^Γ^ is the submatrix of A^Γ^ corresponding to the genotyped individuals, G = (1−α) (ω + ω_b_ x G_V_) + α A ^Γ^, with G_V_ the genomic relationship matrix estimated from the SNPs and scaled following the first scaling method of Van Raden (2008), α a scaling parameter (here equal to 0.05), and ω_a_ and ω_b_ chosen to equate the average inbreeding and the average relationships in G_V_ and A ^Γ^ as in Christensen et al. (2012). As the sample size was relatively small, the matrix H^−1^ was obtained by simply inverting H with the function *solve* (R package *base*).

### Population structure

In order to control for population structure and avoid spurious associations, we computed a distance matrix 1 − H*, where H* is the correlation matrix obtained from H using the function *cov2cor* (R package *stats*), and the minus sign is to convert similarities to distances. We then used Multidimensional Scaling (MDS) on the distance matrix with the function *cmdscale_lanczos* (R package *refund* v0.1; Goldsmith et al., 2022; Miller, 2022) with the parameter ‘k’ set to 5 (hence explaining > 90% of the variance), and ‘eig’ set to ‘TRUE’. Negative eigenvalues were set to null, and the variance explained by an axis was computed as the ratio between its squared eigenvalue and the sum of the squared eigenvalues. Additionally, to assess how well the family structure was captured, we measured the variable V_F_ defined as the average within family variance on the first five axes (the lower the V_F_, the better the family structure is captured). To make each MDS derived from the different subsets comparable with each other, we normalized V_F_ by the sum of squares of the first five eigenvalues.

### Breeding value inferences

The spatially adjusted phenotypes were then used to infer the breeding values with a multitrait single-step Genomic Best Linear Unbiased Predictor (ssGBLUP), as follows:

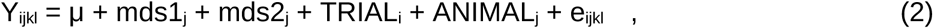

where ‘Y_ijkl_’ is the l-th phenotype of the k-th replicate of the j-th genotype in the i-th trial, μ is the grand mean, ‘mds1_j_’ and ‘mds2_j_’ the fixed effects on the j-th genotype of the first and second axes of the MDS, respectively, ‘TRIAL_i_’ the random effect of the i-th trial, ‘ANIMAL_j_’ the polygenic effect of the j-th genotype, and ‘e_ijkl_’ the residuals. Both the trial effect and the residuals follow a Gaussian distribution with diagonal covariance matrices, and the polygenic effect follows a Gaussian distribution with a covariance matrix equal to k.H x Σ, where k is a scaling parameter equal to 1 - γ/2 (Legarra et al., 2015), H the combined relationship matrix, x the Kronecker product, and Σ the 2×2 covariance matrix between height and vulnerability to rust. Estimates of coefficients and variance components of model (2) were obtained with the function *remlf90* (R package *breedR* v0.12) through one run with the ‘method’ parameter set to ‘em’ until convergence, then another run with the parameter ‘method’ set to ‘ai’ and the parameter ‘progsf90.option’ set to ‘maxrounds 1’. The output estimate of ‘ANIMAL_j_’ is the Estimated Breeding Value (EBV) of the j-th genotype. The additive variance was obtained from the diagonal elements of Σ, and used to estimate heritability (the denominator of which also comprises the residual variance). The predictive ability (PA) of the model was defined as Pearson’s correlation coefficient between the breeding values re-estimated from the effect sizes (see below) and the adjusted phenotypes corrected by the trial effect (i.e., Y_ijkl_ − TRIAL_i_).

### Genome-Wide Association Study (GWAS)

Following the equations of Aguilar et al. (2019), the effect size of the SNPs was back-solved from the EBVs, along with their standard error and their p-values. This process was performed trait by trait (i.e., without accounting for the multitrait dimension) without loss of information, and taking all effects as fixed, as the Pearson’s correlation between the ssGBLUP’s EBV and the breeding values re-estimated by the effect sizes was greater than 0.999.

With the estimated effect sizes and the standard errors, we also computed an alternative to the p-value accounting for multiple testing: the local false sign rate (lfsr) estimated with an empirical Bayes approach for adaptive shrinkage using the function *ash* (R package *ashr* v2.2; Stephens, 2017). In order to detect GWAS peaks (regions in the chromosomes with a high density of significant SNPs), we used a custom peak detection script on the profile of lfsr (Tiret & Milesi, 2021) with default parameters. The script filtered out significant hits that are not statistically detected as a peak which is defined here as a statistically improbable concentration of significant SNPs in a small chromosomic area.

### Neighborhood

In order to account for the effect of the neighborhood, we estimated the breeding values with an additional fixed effect, the local relatedness, defined as the average relationship (from the matrix H) between a focal individual and its neighbors (eight or less if on boundaries; as the king’s moves on a chessboard). Local relatedness was orthogonal to any micro-environmental effect, because of the random block design. The re-estimated breeding values were denoted nEBV. To assess the goodness of fit of the different models, we assessed the Akaike Information Criterion (AIC) with the function *AIC* of the R package *stats*. To assess the extent to which local relatedness enriched phenotypic covariance, we compared the phenotypic covariance between random pairs of neighboring individuals with random pairs from the most related neighbors (highest quartile of local relatedness), sampling for both calculation 10,000 pairs.

### SNP subsets

We investigated different subsets of SNPs, each reflecting different aspects of past selection or demography. We then computed (genomic) local relatedness with the different subsets, so to assess which aspect of the past history is responsible for the association between local relatedness and focal individuals’ phenotype. We selected SNP subsets based on Minor Allele Frequency (MAF), Ancestry Informativeness Coefficient (AIM; Rosenberg et al., 2003), and for each trait, significance in the GWAS, and additive variance. For each feature, we kept one-tenth of the whole set (713 SNPs, a sample size close to previous studies on weighted GBLUP, e.g., Li et al., 2018) with either the smallest or the largest values, ending up with 12 subsets. High MAF, low AIM, high significance in the GWASs, and high additive variance were interpreted as reflecting past selection, and the opposite as reflecting drift and neutral demographic dynamics. AIM was estimated per family with the R script provided by Cappa et al. (2022). Additive variance explained by a SNP was estimated as 2p(1-p)a^2^, where p is the MAF, and a the effect size (assuming no dominance nor interaction deviation of the genetic variance; p.129, Falconer, 1989). We re-estimated the combined relationship matrix for each SNP subset by re-estimating in the equation (1) the genomic relationship matrix G_V_ with the SNP subset. With the re-estimated matrices H, we re-performed the MDSs, and re-estimated the PAs of the EBVs and nEBVs (where local relatedness were measured with the re-estimated H).

### Statistical tests

For each SNP subset, we computed the confidence interval of the Predictive Ability (PA) with a 1000 iteration bootstrap (re-sampling individuals), without re-estimating the covariance matrix Σ. Here, the bootstrap was only used to assess the variance, so that we centered the bootstrapped PA around the true PA. To assess the effect of sampling, these subsets were compared to 1000 random samples of 713 SNPs (the size of the SNP subsets). Comparisons were performed with a Student’s one-sample *t-*test (denoted *t_1_* with a degree of freedom or df of 999 corresponding to the number of bootstrap iterations), or a Welch two-sample *t-*test (denoted *t_2_* with a varying df), both using the function *t.test* (R package *stats*). The sign of the *t* statistic is arbitrarily reported as positive when greater than a baseline (e.g., the true PA), and negative when not. Models were compared with a likelihood ratio test with a df of 1 using the function *lrtest* (R package *lmtest* v0.9; Zeileis & Hothorn, 2002). All scripts were written in R v4.1.3 (R Core Team, 2022).

## Results

Tree height and vulnerability to rust both exhibited high heritabilities (0.469 and 0.666, respectively), high predictive abilities (0.677 and 0.755 respectively), and a strong independence, with an estimated genetic correlation of 0.022. We validated that GWAS peaks were likely not artefactual by swapping phenotypes (permutation test) among family members of the same block (i.e., only randomly mismatching Mendelian sampling): the permutation significantly reduced the average significance of the SNPs (t_1_ < -2.59 × 10^2^, p < 0.001). Only one genetic variant was significantly associated with vulnerability to rust, located on chromosome 1, and none for tree height (Fig. 1A, 1B). The Q-Q plot on vulnerability to rust revealed that some SNPs were significantly associated with a level of significance well above the 95% confidence interval (Fig. 1D); most of them being in the identified peak. None of the SNPs were significantly associated with tree height after Bayesian shrinkage, and Q-Q plot revealed a classic population structure overcorrection (Fig. 1C).

**Figure 1.**
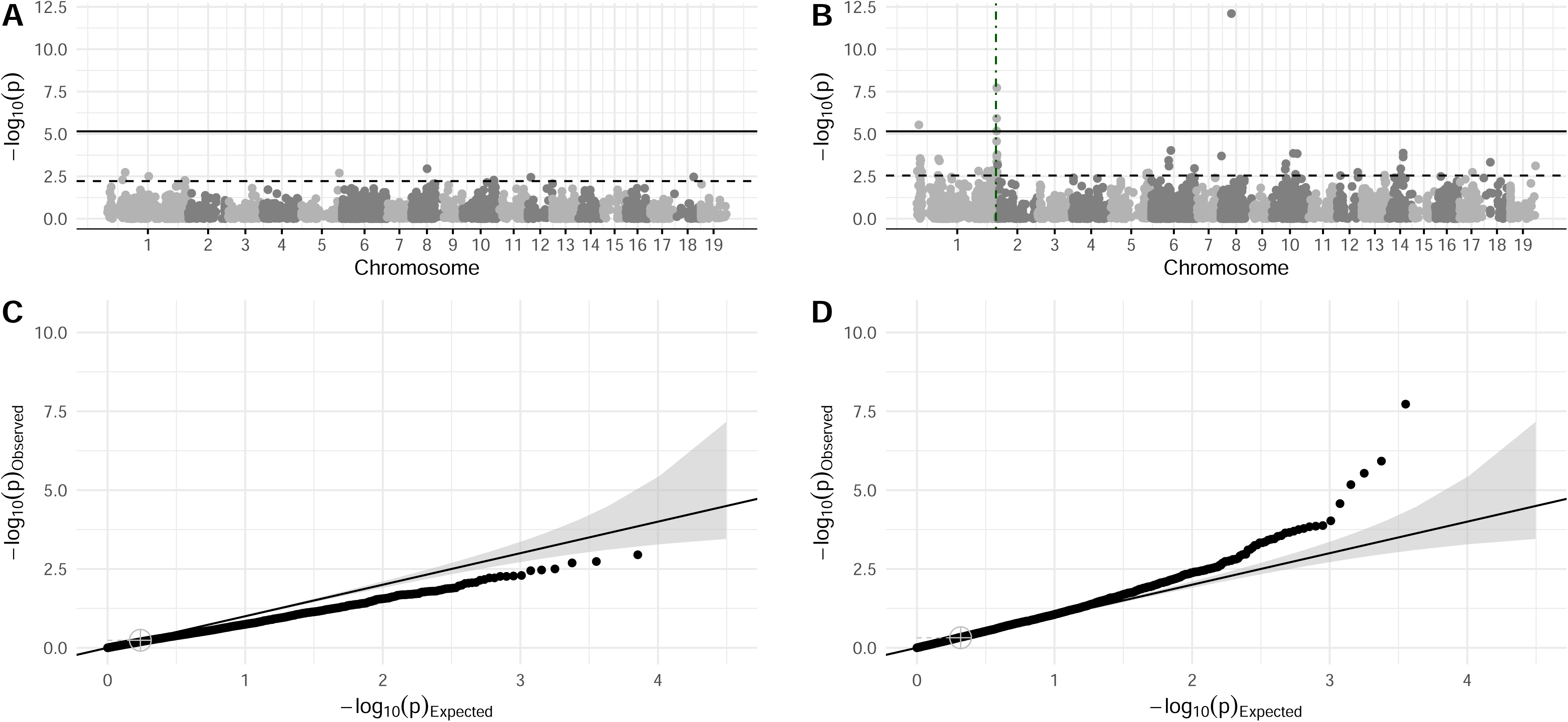
GWAS on tree height and vulnerability to rust in Populus nigra. **(A)**. Manhattan plot of the GWAS on tree height. The dots are the log significance of the markers. The solid horizontal line stands for the 5% significance level, and the dashed horizontal line stands for the 5% FDR level. The vertical dashed lines are position where significant hits were detected as being part of a peak. **(B)**. Idem as panel A, for vulnerability to rust. **(C)**. Q-Q plot of the GWAS on tree height. The dots were the quantile of the markers; the solid line is the 1-slope segment; the gray cross is the position of the median; and the gray area is the 95% confidence interval. **(D)**. Idem as panel C, for vulnerability to rust.

### Local relatedness improves ssGBLUP fit for vulnerability to rust

Although the growth of focal individuals was not affected by their neighbors, damage due to rust was more severe the more related the focal individuals were with their neighbors. Indeed, the genomic models incorporating local relatedness as a covariate had significantly higher likelihood than without for vulnerability to rust (χ^2^ = 9.83, p = 0.002), but not for tree height (χ^2^ = 0.09, p = 0.236). When computing relatedness with the different subsets of SNPs, the Akaike Information Criterion (AIC) decreased by 0.12 for height (when excluding the smallest MAF) and by 7.95 for vulnerability to rust (Fig. S2), suggesting a better fit when including local relatedness in the model for vulnerability to rust. However, for vulnerability to rust, only the subset of SNPs with the largest MAF significantly increased the likelihood of the model (χ^2^ = 4.02, p = 0.04). Likelihood was not significantly improved by SNPs with the largest AIM (χ^2^ = 0.15, p = 0.703) or the largest additive variance (χ^2^ = 2.65, p = 0.104, respectively), and was significantly worse with other subsets (χ^2^ > 6.66, p < 0.010). For tree height, no subset of SNPs improved the likelihood (χ^2^ < 3.48, p > 0.062). In other words, focal individuals were more affected by rust when they had related individuals in their vicinity. The covariance between random pairs of neighboring individuals’ vulnerability to rust compared to the most related neighbors’ pairs increased from an average covariance of 0.28 to 0.43, showcasing a systematic association despite the randomized block design.

In spite of genomic models being sensitive to local relatedness, improvement of predictive abilities (PA) was limited (Fig. S3; see Supplementary materials for further details) - probably due to a lack of power. Indeed, incorporating local relatedness did not significantly change the PA neither with the whole set of SNPs (|t_2_| < 0.594, p > 0.553), nor with any of the alternative subsets (|t_2_| < 1.03, p > 0.301), with a slightly better, but non-significant PA when incorporating local relatedness for SNPs with the smallest MAF on vulnerability to rust (t_2_ = 1.73, df = 2.00 × 10^3^, p = 0.0841), and SNPs with the smallest tree height significance on vulnerability to rust (t_2_ = 1.97, df = 2.00 × 10^3^, p = 0.0491).

### The impact of local relatedness on vulnerability to rust is <u>likely</u> due to past family selection

The objective here is to assess which SNP subset reveals which aspect of the past history (selection or drift), knowing that local relatedness is mainly associated to vulnerability to rust through SNPs with large MAF. The population under study is structured in a hierarchical way: first into populations, then into families. Population structure, captured by the first eigenvalues (Patterson et al., 2006), substantially shaped genetic diversity, as the first two eigenvalues alone explained more than 50% of the variance (40.0% and 24.5% respectively; Fig. S1). Because the population underwent family selection during breeding in the past, family structure is strong and can even be apprehended visually (Fig. 2). Several statistics significantly supports the strong family structure: (i) randomly shuffling the family ID increased by 9.96 times the within “family” variance (t_1_ = 3.70 × 10^3^, p < 0.001); (ii) when restricting to common alleles, the first two eigenvalues of the PCA still explained a large part of the variance (56.2%), showing that family structure is bound to selection, as expected under family selection.

**Figure 2.**
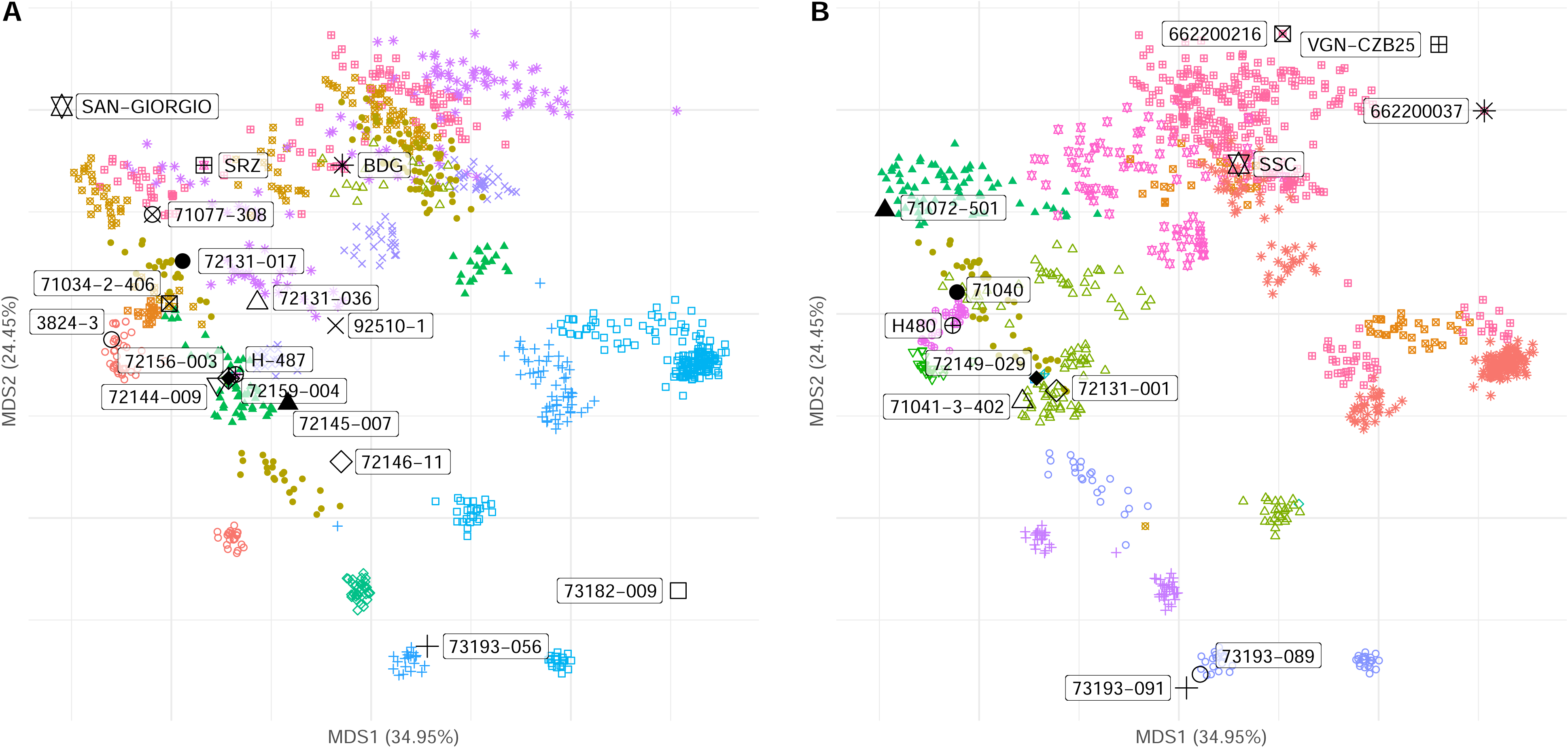
The first two axes of the Multidimensional Scaling (MDS) of Populus nigra’s genotyping data. **(A)**. MDS with labels of the fathers (with black large shapes). Colored and smaller shapes are the offspring, the shape corresponding to its father. **(B)**. The same MDS as in panel A., but labelled for the mothers.

In the case of the investigated populations, family structure can reveal neutral past dynamics, or past family selection. In order to understand what drove the family structure, similarly to using annotation to focus on some SNPs, we subset 10% of the SNPs first randomly and then according to their MAF, AIM, significance in a GWAS, or additive variance, thus redefining relatedness with a different set of linkage disequilibrium. In our dataset, random samples of SNPs, performed quite poorly compared to the whole set of SNPs (t_1_ = 2.12 × 10^2^, p < 0.001). However, SNPs with the largest AIM, performing better than random samples (t_1_ = 3.86 × 10^2^, p < 0.001; Fig. 3), and even than the whole set; and SNPs with the largest MAF and the largest additive variance, performing better than random samples (t_1_ > 1.09 × 10^2^, p < 0.001). The improvement in goodness of fit due to incorporating local relatedness in the genomic models is thus due to any forces structuring the family, likely past family selection since it increases the MAF.

**Figure 3.**
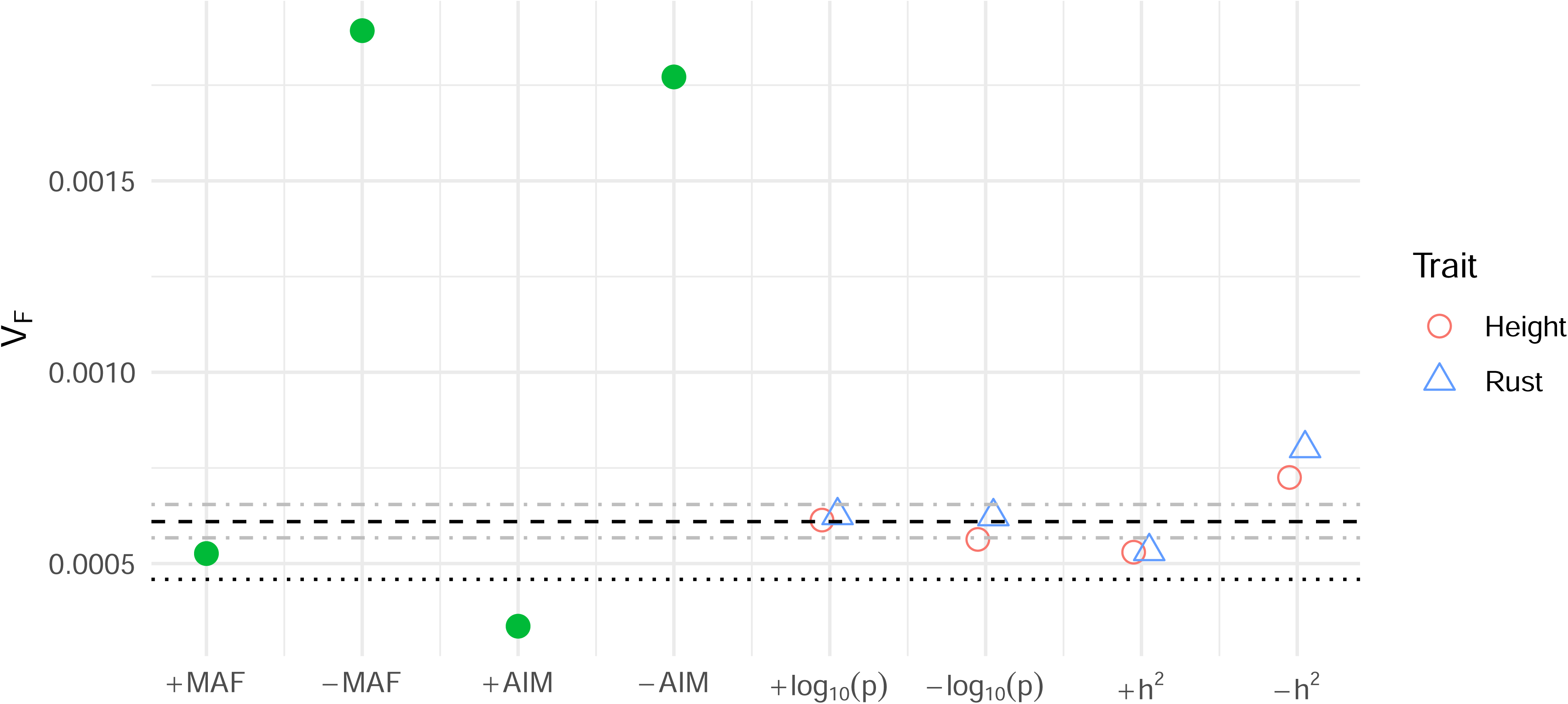
Within family variance (V_F_) for each subset of SNPs. The black dashed horizontal line is the average V_F_ of randomly sampled SNPs, the gray dashed horizontal lines delimit the 95% confidence interval, and the black horizontal dotted line is V_F_ of the entire set of SNPs. Red dots are when selected for tree height, and blue for vulnerability to rust (green dots are when this distinction is not relevant). SNP subsets: largest or smallest MAF (+ or - MAF), largest or smallest AIM (+ or - AIM), most or least significant (+ or - log_10_(p)), and most or least heritable for tree height (+ or - h^2^).

## Discussion

The success of prediction based on genomic models relies on the ability of genotypic data to capture the covariance between candidate loci, thereby reflecting how much genetic similarity explains phenotypic similarities (Powell et al., 2010). An additional source of phenotypic covariances are micro-environmental effects, and accounting for spatial autocorrelation can enhance genomic predictions (Cappa et al, 2022). In our case, on top of spatial autocorrelation, we accounted for local relatedness, another source of heterogeneity. The significant increase of the likelihood when adding local relatedness as a covariate shows that part of the plant-to-plant interaction can be ascribed to the relatedness with neighbors, at least for vulnerability to rust.

### Different genetic architectures and local relatedness

Height is known to have a highly polygenic genetic architecture (Du et al., 2016), and resistance to rust to have a mono- or oligogenic architecture (e.g., Jorge et al., 2005), implying that relatedness – as the amount of shared alleles – may have different consequences on those two traits: genetic redundancy (Nowak et al., 1997; Láruson et al., 2020) of height probably making it less sensitive to local relatedness than resistance to rust. The low genetic correlation between both traits is consistent with some previous reports (e.g., Beaulieu et al., 2020; Liu et al., 2022), though not all (e.g., Lenz et al., 2021; Cappa et al., 2022); we will assume in our case that orthogonality is strong enough to show that the traits are non-redundant, with independent architectures.

The differences in genetic architecture between the two traits studied, as well as the design of the SNP array, may have determined the results in terms of association, with only one major gene detected (as a peak) for rust, in contrast to the absence of associations for the more polygenic trait. Indeed, a major caveat of our analysis is that the SNP chip array we used is enriched for QTLs and candidate expression genes related to resistance to rust during its development (Faivre-Rampant et al., 2016), but not for height. While there may be a risk of ascertainment bias due to the frequency criteria used and the design of the array, this array was constructed using a very large diversity panel that is representative of the western distribution of the species. Several studies have compared different technologies (e.g., SNP arrays, genotyping-by-sequencing, etc.) and found that ascertainment bias was negligible for monitoring diversity trends and estimating relatedness (Torkamaneh et al., 2015; Negro et al., 2019). Although biases may exist, the effects are unlikely to be large enough to invalidate the results. However, further investigation with alternative sequencing technologies would be needed to validate this conclusion. Another element that may have influenced the association for rust is its phenotypic scoring, which is based on a qualitative scale of nine grades, which might artificially grade an inoculum pressure linearly. Overall, even if the results should be taken cautiously, it is not uncommon to encounter a simpler genetic architecture for response to biotic constraints compared to complex quantitative traits (e.g., gene-for-gene resistance; Flor, 1971; Jorge et al., 2005). Consequently, many haplotype combinations can lead to tall individuals, but few to resistance to rust, because a polygenic architecture is redundant; in other words, when individuals are related, their probability of both being resistant is higher than that of both being tall (individuals sharing rare alleles are closer than when sharing common alleles; Van Raden, 2008). This reasoning is specifically adapted in the case of factorial crossing where all possible crosses have been performed numerous times. Therefore, local relatedness would increase with a vulnerability to rust that is shared within the neighborhood, even if it is not inducing vulnerability to rust *per se*.

### Is local relatedness the cause of local phenotypic similarities?

Interactions between neighbors may depend on local relatedness, and whether this interaction is positive or negative depends on the relative strength of kin selection (Hamilton, 1964) and niche partitioning (Silvertown, 2004). As mentioned above, the importance of group selection in nature is still debated, but local relatedness has been shown to be perceived through kin recognition in plant species (e.g., Dudley et al., 2007; Murphy et al., 2009). Such recognition could then lead to favorable (kin selection) or unfavorable (niche partitioning) interactions. One such detrimental interaction is competition, which can be accounted for in a quantitative genetic model in the DGE/IGE framework (Griffing, 1967; potential applications in Muir, 2005). Competition occurs when DGE and IGE are negatively correlated. A positive correlation from such a model would suggest a cooperative synergistic interaction. Although adding complexity to the model, including competition has been shown to reduce bias in covariance estimation (Costa e Silva et al., 2013). In our case, in addition to attempts to include interactions, IGE resulted in a lower predictive ability than with the ssGBLUP baseline (results not shown).

Once it has been shown that neighbourhood relatedness is indeed correlated with similarity at focal phenotypes, the difficulty in disentangling kin selection from niche partitioning is to determine whether the relatedness is a cause, as an expression of kin selection or niche competition, or a co-product, as an expression of a confounding between relatedness and interaction through phenotypes, where the focal phenotype would be modified by that of its neighbors through plant-to-plant interactions. If phenotypic interaction increases when phenotypes are similar (e.g., competition for resource uptake), it can be confused with the effect of relatedness, as genetically related individuals tend to have similar phenotypes. However, if local relatedness was only a proxy for highly related resistant to rust neighborhoods, average vulnerability to rust would increase with local relatedness but not covariance of vulnerability to rust (i.e., a systematic and coherent increase in vulnerability to rust), because of the randomized block design.

The results hence suggest that shared vulnerability is likely the cause of rust damage amplification in the related neighborhood. The amplifying effect might be even stronger in perennials as shared vulnerability can be perceived by successive generations of pathogens. This phenomenon is consistent with the increased disease resilience conferred by varietal association in a field (e.g., Smithson and Lenne, 1996; Burdon and Thrall, 2009), or the detrimental effect of lack of diversity in several plant species (Zhu et al., 2000; Liu et al., 2020), but would need further investigations. On the other hand, the fact that family structure was mainly resulting from allele identity sharing at SNPs likely associated with selection (following the hypothesis of Yang et al., 2010 and Biddanda et al., 2020) suggest that relatives were co-selected (relatedness as a confounding effect), as is often the case in breeding programs implementing family selection. An inevitable consequence is that, as seen in crop selection, interesting genotypes for kin selection/cooperation have been lost during domestication (Fréville et al., 2022), making it difficult for genomic selection to account for local relatedness (as predictive abilities show).

## Conclusion

The effect of the relatedness of a focal individual with its neighbors was investigated on tree height and vulnerability to rust, with 10,301 trees of 1,452 genotypes across four trials. The results show that incorporating local relatedness as an additional factor in GBLUP models has a significantly greater influence on estimation of resistance to rust than on tree height, though its overall effect on genomic predictions was limited. Whether there was kin selection/competition in addition to the phenotypic interaction, or whether we simply lacked statistical power, is difficult to assess. Selection on local relatedness due to kin selection or niche partitioning - selection on relatedness (favoring specific plant-to-plant interaction due to relatedness) - might be confused with the fact that selection leads to phenotypic convergence, that in turn leads to a genotypic convergence - a co-selection of related (selecting for phenotypes that accidentally but inevitably selected for related individuals). Further investigation is required to elucidate the relationship between genetic architecture and the impact of relatedness on phenotypes.

## Acknowledgements

The authors acknowledge the Uppsala University and the European Union’s Horizon 2020 B4EST for basic functioning and postdoctoral grant for MT. The authors would like to thank Véronique Jorge and Remy Gobin of the BioForA unit (INRAE, ONF, Orléans, France) for their help in compiling the data. The computations and data handling were enabled by resources provided by the Swedish National Infrastructure for Computing (SNIC 2017-7-296) at UppMax partially funded by the Swedish Research Council through grant agreement no. 2018-05973.

## Author Contributions

MT was responsible for writing the report, conducting the search, extracting and analysing data, interpreting results, updating reference lists. LS was responsible for designing the experiments and screening potentially eligible studies. ML and LS contributed to interpreting results and provided feedback on the report.

## Competing Interests

The authors declare that there is no conflict of interest.

## Data Archiving

The data that support the findings of this study are openly available on DATA INRAE at https://data.inrae.fr/privateurl.xhtml?token=b79ab1ca-ebb9-47c6-9272-1568c0c33d70.

## Notes

https://data.inrae.fr/privateurl.xhtml?token=b79ab1ca-ebb9-47c6-9272-1568c0c33d70.

### Competing Interest Statement

The authors have declared no competing interest.

### Summary of Updates

Clarification in the abstract, introduction, results and discussion. Nuances on the scope of the results.

