## Supplementary material for "The role of relatedness within a neighborhood in plant-to-plant interaction: effect on height growth and rust damage in *Populus nigra*"

### Supplementary materials

##### Tables

| Trial | 1999-2000 | 2012-2013 | 2014-2016 | 2017-2018 | **Total** |
| --- | --- | --- | --- | --- | --- |
| # full-sib families | 14 | 9 | 16 | 27 | **42** |
| # half-sib families | 1 | 1 | 5 | 1 | **5** |
| # genotypes | 517 (14) | 273 (10) | 320 (79) | 411 (15) | **1,452 (105)** |
| # blocks | 8 | 6 | 6 | 6 | **26** |
| # trees | 4,224 | 1,656 | 1,920 | 2,501 | **10,301** |

**Table S1.** Composition of the four experimental trials. # *x* = number of *x*. The number of genotypes in the Unknown Parent Group is indicated within parentheses.

|  | Tree height | | | | Rust vulnerability | | | |
| --- | --- | --- | --- | --- | --- | --- | --- | --- |
|  | Correlation | | Peak | | Correlation | | Peak | |
|  | EBV | nEBV | EBV | nEBV | EBV | nEBV | EBV | nEBV |
| - MAF | 0.829 | 0.829 | - | - | 0.762 | 0.762 | 49,917,223 | 49,903,967 |
| - MAF | 0.694 | 0.694 | - | - | 0.558 | 0.558 | 39,658,416 | 42,300,533 |
| - AIM | 0.842 | 0.842 | - | - | 0.764 | 0.764 | 47,158,157 | 47,158,157 |
| - AIM | 0.791 | 0.791 | - | - | 0.745 | 0.745 | 49,503,130 | 49,782,094 |
| + log_10_(p): height | 0.595 | 0.595 | - | - | 0.742 | 0.741 | - | - |
| - log_10_(p): height | 0.875 | 0.875 | - | - | 0.763 | 0.763 | - | 49,903,967 |
| + log_10_(p): rust | 0.812 | 0.812 | - | - | 0.486 | 0.486 | 47,128,425 | 47,158,157 |
| - log_10_(p): rust | 0.826 | 0.826 | - | - | 0.915 | 0.915 | 49,393,096 | 47,158,157 |
| + h^2^: height | 0.584 | 0.584 | - | - | 0.710 | 0.710 | 42,485,193 | 42,485,193 |
| - h^2^: height | 0.869 | 0.869 | - | - | 0.763 | 0.763 | 2,133,128 | 2,133,128 |
| + h^2^: rust | 0.786 | 0.786 | - | - | 0.472 | 0.472 | 47,158,157 | 47,158,157 |
| - h^2^: rust | 0.835 | 0.835 | - | - | 0.902 | 0.902 | 49,404,715 | 49,403,968 |

**Table S2.** Correlation of effect sizes between SNP subsets and the entire set of SNPs, and peak positions, for tree height and rust vulnerability. GWAS on subsets of SNPs resulted with effect sizes with intermediate to high correlation with the true effect sizes, and peaks were mostly located at the same positions. SNP subsets: largest or largest MAF (+ or - MAF), largest or smallest AIM (+ or - AIM), most or least significant for tree height (+ or - log_10_(p): height), most or least significant for rust vulnerability (+ or - log_10_(p): rust), most or least heritable for tree height (+ or - h^2^: height), and most or least heritable for tree height (+ or - h^2^: rust). EBV: Estimated Breeding Value; nEBV: Estimated Breeding Value including local relatedness.

##### Figures

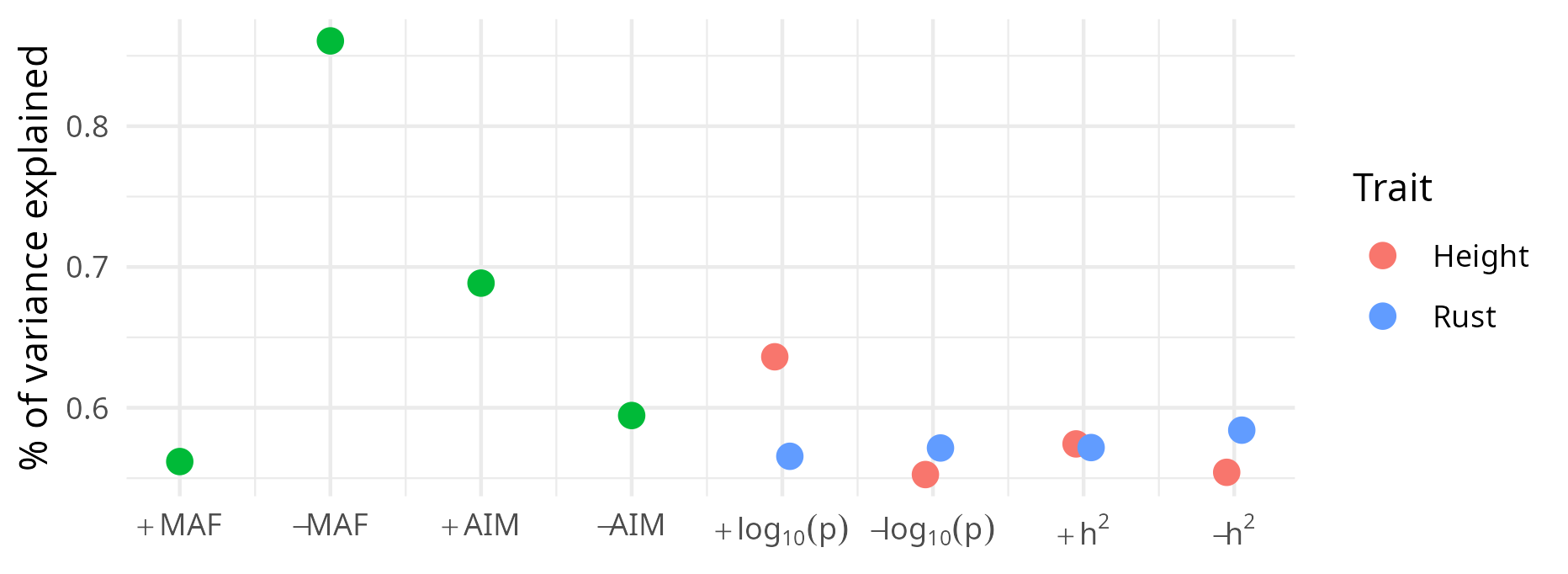

**Figure S1.** Proportion of variance explained by the first two axes of the Multidimensional Scaling (MDS) of the genotypes. Red dots are when selected for tree height, and blue for rust vulnerability. SNP subsets: largest or largest MAF (+ or - MAF), largest or smallest AIM (+ or - AIM), most or least significant (+ or - log_10_(p)), and most or least heritable for tree height (+ or - h^2^).

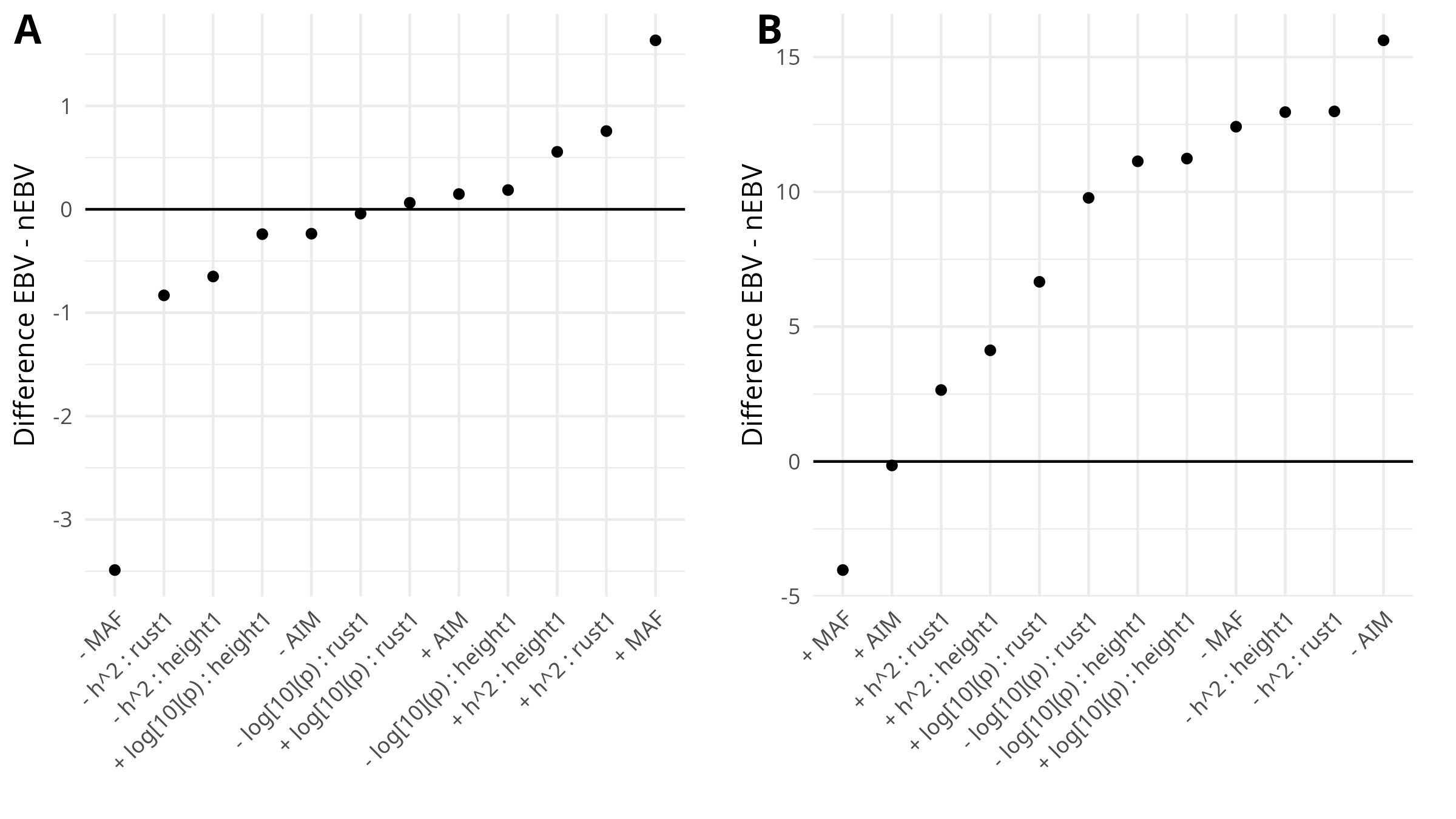

**Figure S2**. Difference of AIC between EBV and nEBV (including local relatedness as a covariate), for different subset of SNPs: largest or largest MAF (+ or - MAF), largest or smallest AIM (+ or - AIM), most or least significant (+ or - log_10_(p)), and most or least heritable for tree height (+ or - h^2^). When the difference is positive, nEBV is a better model than EBV. **(A)**. Tree height. **(B)**. Rust vulnerability.

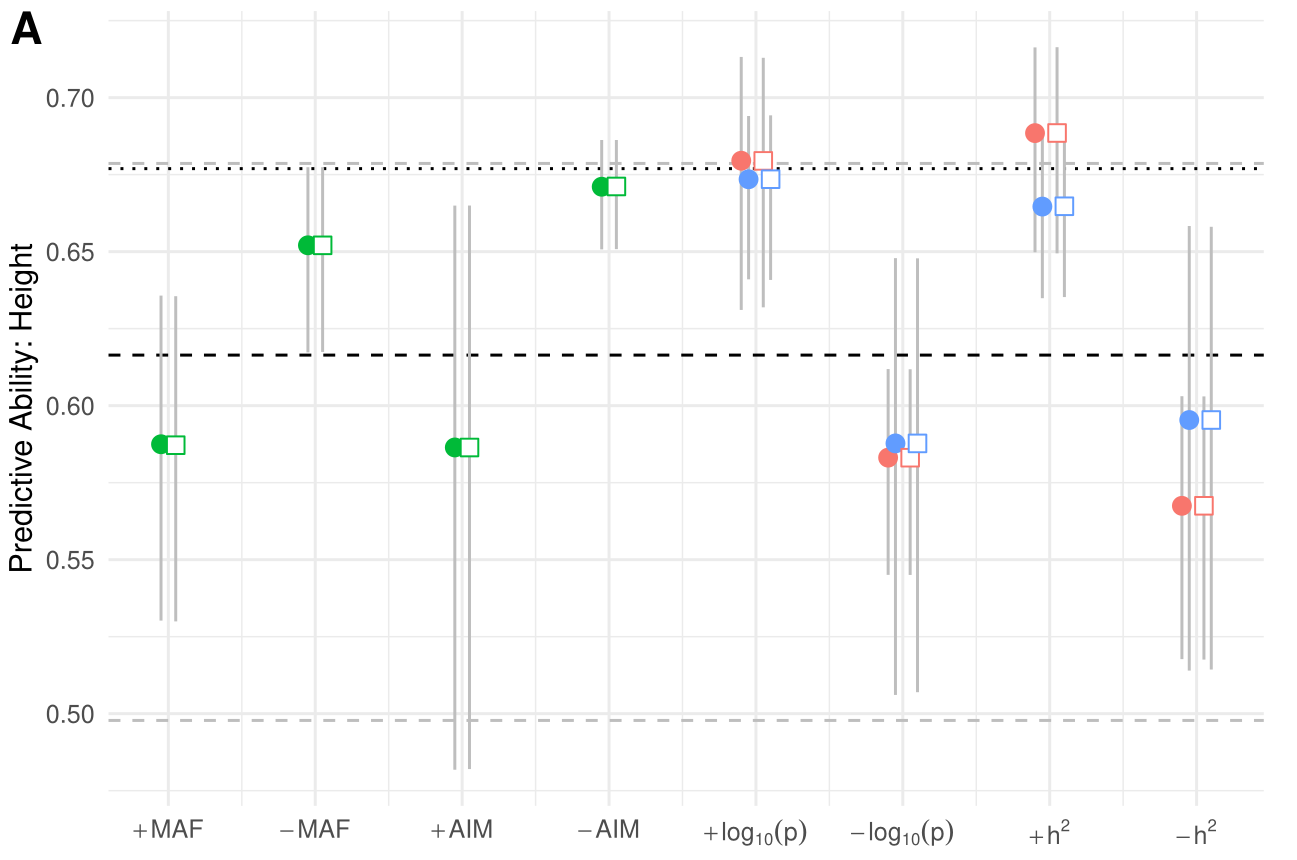

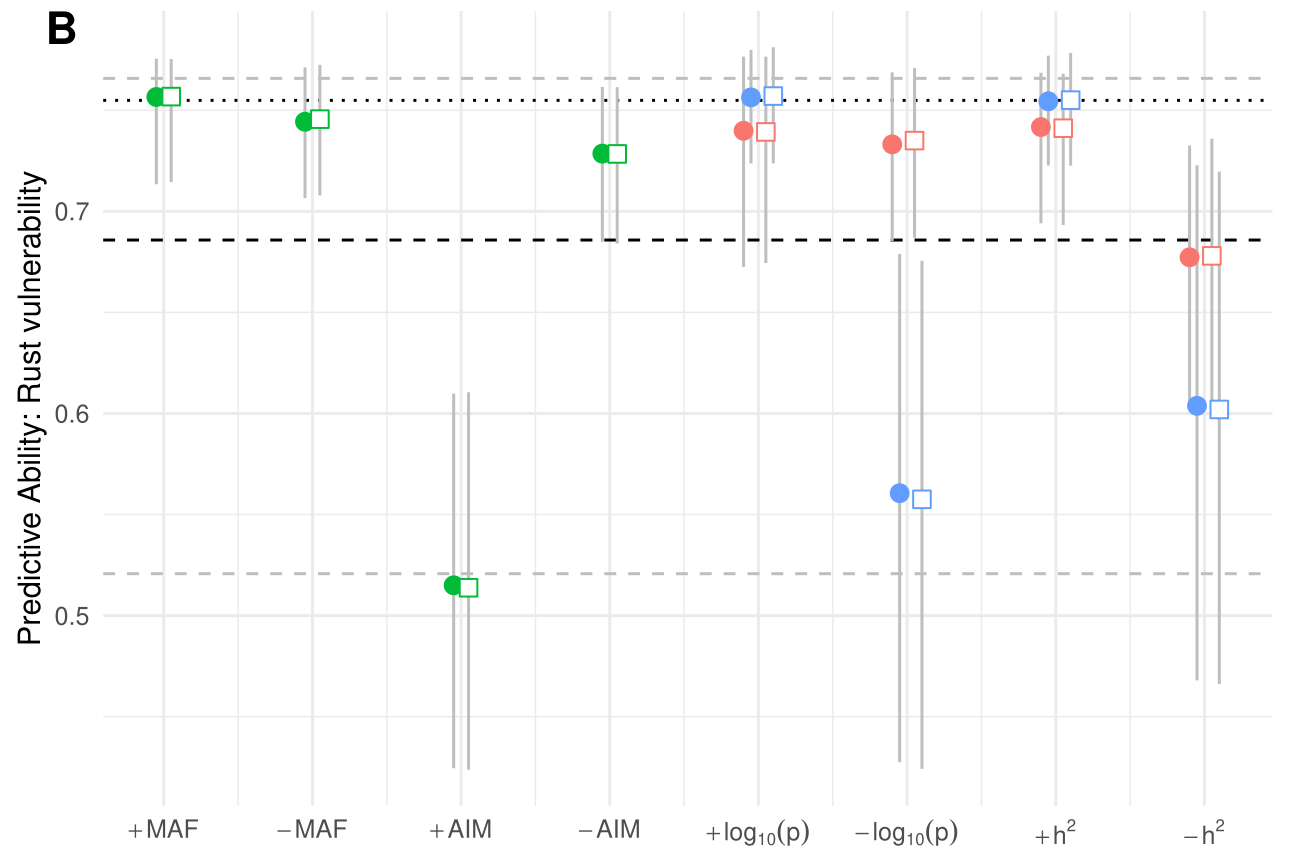

**Figure S3.** Predictive ability (PA) of different genomic models. **(A)**. PA for tree height, for different SNP subsets (along the x-axis). The black dotted horizontal line is the PA of the entire set of SNPs, the black dashed horizontal line is the average PA of randomly sampled SNPs, and the gray dashed horizontal lines delimit the 95% confidence interval. Cercles are when models do not include local relatedness as a covariate, and squares when they do. Gray vertical lines display the 95% confidence interval computed from 1000 iteration bootstraps. Red shapes are when selected for tree height, and blue for rust vulnerability. SNP subsets: largest or largest MAF (+ or - MAF), largest or smallest AIM (+ or - AIM), most or least significant (+ or - log_10_(p)), and most or least heritable for tree height (+ or - h^2^). **(B)**. Idem as panel A, for rust vulnerability.

#### Predictive abilities were exclusively driven by the relationship matrix

The different definitions of the relationship matrix led to different predictive abilities (PA), depending on the combination of traits and SNP subsets (Fig. S3). PA for the whole set of SNPs was significantly higher than for random samples for both tree height (t_1_ = 3.97 x 10, p < 0.001) and rust vulnerability (t_1_ = 3.31 x 10, p < 0.001). However, some random samples had a higher PA than the whole set, and the amount of such samples was higher for rust vulnerability (1.20%) than for tree height (0.39%). This difference is probably due to the SNP chip design (even random SNPs are likely QTLs for rust vulnerability). The most significant SNPs for rust vulnerability reached a PA significantly higher than that of random samples (t_2_ = 3.30 x 10, df = 1.09 x 10^3^, p < 0.001), than the least significant SNPs (t_2_ = 9.27 x 10, df = 1.09 x 10^3^, p < 0.001), and even than the whole set (t_1_ = 3.27, p = 1.1 x 10^-3^). Likewise, for tree height, the most significant SNPs reached a PA significantly higher than that of random samples (t_2_ = 3.79 x 10, df = 1.37 x 10^3^, p < 0.001), than the least significant SNPs (t_2_ = 1.11 x 10^2^, df = 1.92 x 10^3^, p < 0.001), and than the whole set (t_1_ = 3.82, p < 0.001).

Although the subsets of SNPs significantly structuring families (largest MAF, AIM, and heritability) were significantly different from random samples in regards to PA (|t_2_| > 1.41 x 10, p < 0.001), most did not reach the level of the whole set (t_2_ < -2.20 x 10, p < 0.001), with the exceptions of SNPs with the largest MAF on rust vulnerability (t_1_ = 3.39, p < 0.001), SNPs with the largest tree height heritability on tree height (t_1_ = 2.14 x 10, p < 0.001), and SNPs with the largest rust vulnerability heritability on rust vulnerability (t_1_ = -1.02, p = 0.31). Family structure was not necessarily overlapping genomic performance in this population. Quite logically, selecting the least significant or the least heritable rust vulnerability SNPs had a stronger impact on rust vulnerability than on tree height with regards to PA, and conversely with the least significant or the least heritable tree height SNPs (Fig. 4). However, the influence of one trait on another was asymmetrical, as SNPs with the smallest significance on rust vulnerability resulted with a PA of tree height only slightly different than SNPs with the smallest significance on tree height (t_2_ = -3.61, df = 1.43 x 10^3^, p < 0.001), but SNPs with the smallest significance on tree height ended up with a significantly higher PA of rust vulnerability than SNPs with the smallest significance on rust vulnerability (t_2_ = 79.61, df = 1.21 x 10^3^, p < 0.001).
